## Supplemental figures and tables for "Comparative RNA-Seq Analyses of *Drosophila* Plasmatocytes Reveal Gene Specific Signatures In Response To Clean Injury And Septic Injury"

A

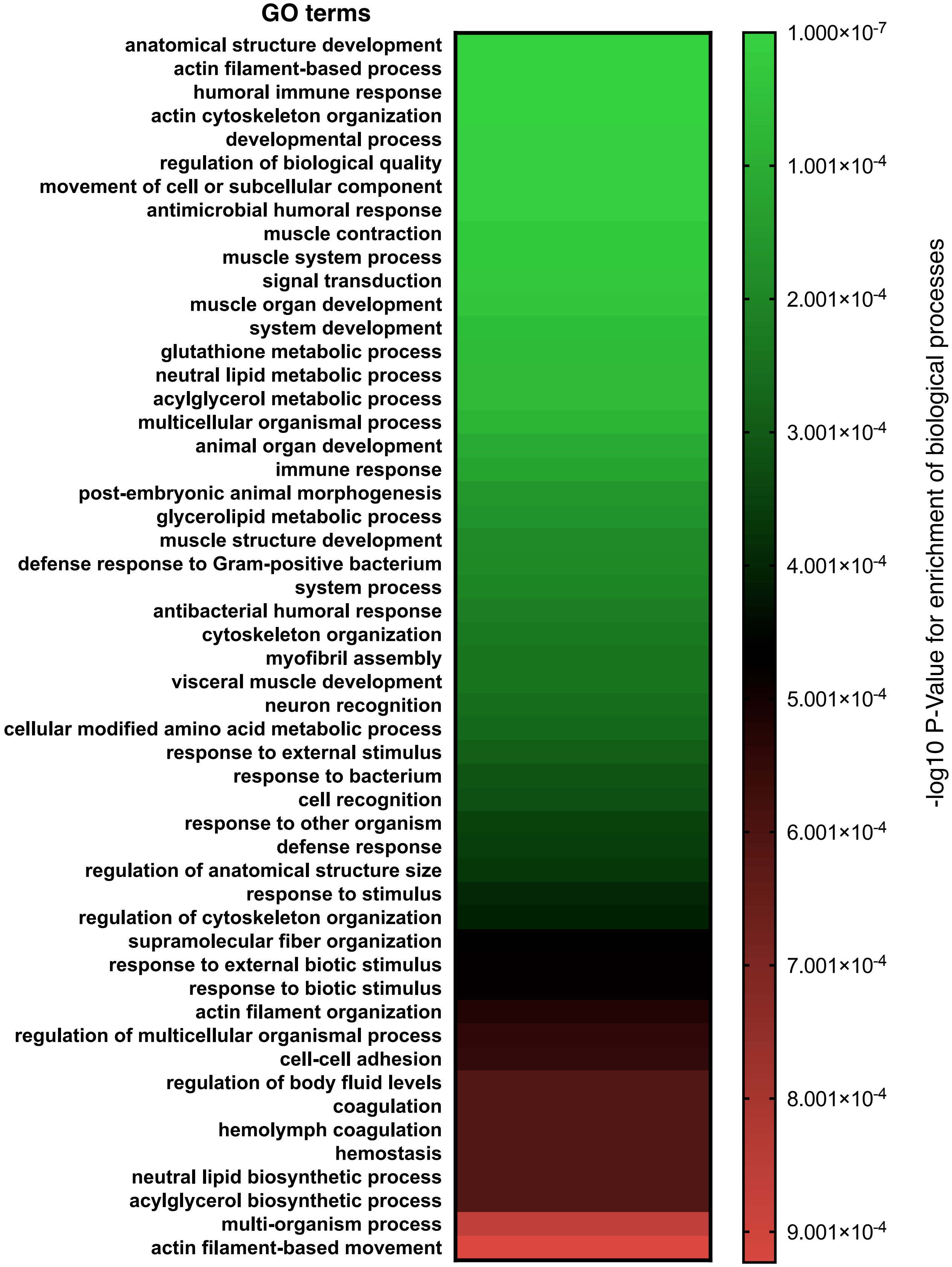

GO terms
anatomical structure development
actin filament-based process
humoral immune response
actin cytoskeleton organization
developmental process
regulation of biological quality
movement of cell or subcellular component
antimicrobial humoral response
muscle contraction
muscle system process
signal transduction
muscle organ development
system development
glutathione metabolic process
neutral lipid metabolic process
acylglycerol metabolic process
multicellular organismal process
animal organ development
immune response
post-embryonic animal morphogenesis
glycerolipid metabolic process
muscle structure development
defense response to Gram-positive bacterium
system process
antibacterial humoral response
cytoskeleton organization
myofibril assembly
visceral muscle development
neuron recognition
cellular modified amino acid metabolic process
response to external stimulus
response to bacterium
cell recognition
response to other organism
defense response
regulation of anatomical structure size
response to stimulus
regulation of cytoskeleton organization
supramolecular fiber organization
response to external biotic stimulus
response to biotic stimulus
actin filament organization
regulation of multicellular organismal process
cell-cell adhesion
regulation of body fluid levels
coagulation
hemolymph coagulation
hemostasis
neutral lipid biosynthetic process
acylglycerol biosynthetic process
multi-organism process
actin filament-based movement
1.000×10-7
1.001×10-4
2.001×10-4
3.001×10-4
4.001×10-4
5.001×10-4
6.001×10-4
7.001×10-4
8.001×10-4
9.001×10-4
-log10 P-Value for enrichment of biological processes


Figure S1

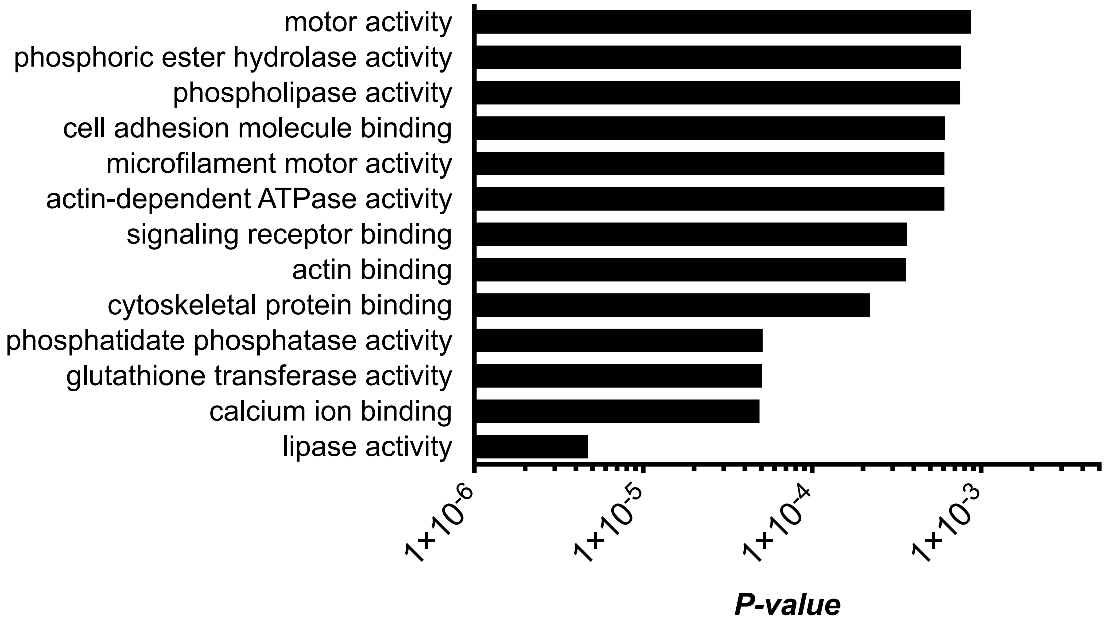
motor activity
phosphoric ester hydrolase activity
phospholipase activity
cell adhesion molecule binding
microfilament motor activity
actin-dependent ATPase activity
signaling receptor binding
actin binding
cytoskeletal protein binding
phosphatidate phosphatase activity
glutathione transferase activity
calcium ion binding
lipase activity
1×10-6
1×10-5
1×10-4
1×10-3
P-value

Figure S2

| CG | Full name | FC |
|---|---|---|
| Extracellular matrix | | |
| Extracellular matrix components | | |
| *CG6953* | *fat-spondin* | 19.10 |
| *CG9280* | *Glutactin* | 18.62 |
| *CG6378* | *Secreted protein, acidic, cysteine-rich* | 16.22 |
| *CG2198* | *Amalgam* | 9.40 |
| *CG3322* | *Laminin B2* | 8.74 |
| *CG7123* | *Laminin B1* | 7.21 |
| *CG16858* | *viking* | 6.40 |
| *CG10236* | *Laminin A* | 5.78 |
| *CG3083* | *Peroxiredoxin 6005* | 6.62 |
| *CG5826* | *Peroxiredoxin* | 3.14 |
| Cytoskeleton organization | | |
| GTPases superfamily | | |
| *CG9366* | *Rho-like* | 5.97 |
| *CG12530* | *Cdc42* | 4.44 |
| *CG8416* | *Rho1* | 4.24 |
| *CG8556* | *Rac2* | 4.18 |
| *CG2248* | *Rac1* | 2.89 |
| Actin binding proteins | | |
| *CG8936* | *Arp2/3 complex, subunit 3B* | 27.62 |
| *CG8978* | *Arp2/3 complex , subunit 1* | 11.92 |
| *CG32858* | *singed* | 9.91 |
| *CG30173* | *Haematopoietic stem/progenitor cell protein 300* | 7.39 |
| *CG5869* | *Glia maturation factor* | 6.54 |
| *CG12363* | *Dynein light chain 90F* | 6.51 |
| *CG9881* | *Arp2/3 complex , subunit 5* | 6.43 |
| *CG15112* | *enabled* | 6.33 |
| *CG9553* | *Chickadee* | 5.80 |
| *CG4636* | *SCAR* | 4.74 |
| *CG10954* | *Arp2/3 complex , subunit 2* | 4.74 |
| *CG9115* | *myotubularin* | 4.27 |
| *CG3265* | *Eb1* | 4.14 |
| *CG4931* | *specifically Rac1-associated protein 1* | 3.90 |
| *CG7558* | *Actin-related protein 3* | 3.88 |
| *CG9749* | *Abelson interacting protein* | 3.72 |
| *CG9901* | *Actin-related protein 2* | 2.82 |
| *CG4560* | *Arp2/3 complex , subunit 3A* | 2.05 |

| CG | Full name | FC |
|---|---|---|
| Membrane trafficking | | |
| Rab GTPases family | | |
| *CG5771* | *Rab11* | 6.35 |
| *CG3870* | *RabX1* | 4.93 |
| *CG9575* | *Rab35* | 3.98 |
| *CG6601* | *Rab6* | 2.97 |
| *CG17060* | *Rab10* | 2.66 |
| *CG8287* | *Rab8* | 2.37 |
| *CG3269* | *Rab2* | 2.19 |
| *CG4921* | *Rab4* | 2.02 |
| Vesicle trafficking | | |
| *CG4764* | *Vacuolar protein sorting 29* | 9.21 |
| *CG10711* | *Vacuolar protein sorting 36* | 4.46 |
| *CG4071* | *Vacuolar protein sorting 20* | 4.14 |
| *CG14804* | *Vacuolar protein sorting 26* | 4.04 |
| *CG6259* | *Vacuolar protein sorting 60* | 3.43 |
| *CG14750* | *Vacuolar protein sorting 25* | 3.06 |
| *CG17828* | *Vacuolar protein sorting 37A* | 2.71 |
| *CG12770* | *Vacuolar protein sorting 28* | 2.13 |
| *CG5625* | *Vacuolar protein sorting 35* | 2.13 |
| *CG7146* | *Vacuolar protein sorting 39* | 2.01 |
